# Frictional effects on RNA folding: Speed limit and Kramers turnover

**DOI:** 10.1101/376558

**Authors:** Naoto Hori, Natalia A. Denesyuk, D. Thirumalai

**Affiliations:** Department of Chemistry, University of Texas, Austin, TX; Biophysics program, Institute for Physical Science and Technology, University of Maryland, College Park, MD

## Abstract

We investigated frictional effects on the folding rates of a human Telomerase hairpin (hTR HP) and H-type pseudoknot from the Beet Western Yellow Virus (BWYV PK) using simulations of the Three Interaction Site (TIS) model for RNA. The heat capacity from TIS model simulations, calculated using temperature replica exchange simulations, reproduces nearly quantitatively the available experimental data for the hTR HP. The corresponding results for BWYV PK serve as predictions. We calculated the folding rates (*k*_F_s) from more than 100 folding trajectories for each value of the solvent viscosity (*η*) at a fixed salt concentration of 200 mM. Using the theoretical estimate (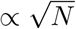 where *N* is number of nucleotides) for folding free energy barrier, *k*_F_ data for both the RNAs are quantitatively fit using one dimensional Kramers’ theory with two parameters specifying the curvatures in the unfolded basin and the barrier top. In the high-friction regime (*η* ≳ 10^−5^ Pa·s), for both HP and PK, *k*_F_s decrease as 1/*η* whereas in the low friction regime *k*_F_s increase as *η* increases, leading to a maximum folding rate at a moderate viscosity (~ 10^−6^ Pa·s), which is the Kramers turnover. From the fits, we find that the speed limit to RNA folding at water viscosity is between (1 − 4)*μ*s, which is in accord with our previous theoretical prediction as well as results from several single molecule experiments. Both the RNA constructs fold by parallel pathways. Surprisingly, we find that the flux through the pathways could be altered by changing solvent viscosity, a prediction that is more easily testable in RNA than proteins.

## Introduction

The effects of friction on barrier crossing events, with a rich history,^1,2^ have also been used to obtain insights into the dynamics and folding of proteins. For example, in a pioneering study, Eaton and coworkers established that accounting for the internal friction is needed to explain experiments in the ligand recombination to the heme in myoglobin.3 Only much later, the importance of internal friction, a concept introduced in the context of polymer physics,4 in controlling the dynamics of folded and unfolded states of proteins has been appreciated in a number of experimental^5–8^ and theoretical^9–13^ studies. Theoretical studies^14,15^ also showed that folding rates of the so-called two-state folders are in accord with the theory of Kramers,^16^ who determined the dependence of the rate of crossing a barrier in one dimension as a function of the viscosity of the surrounding solvent. The timeless Kramers’ theory showed that the rate should increase linearly with *η* at small *η* and decrease as 1/*η* at large *η*. The change from small *η* behavior to 1/*η* dependence with a maximum at intermediate viscosity values is often referred to as the Kramers turnover.^1,2,17^

Kramers’ theory has been used to understand frictional effects of the solvent in various reactions, from diffusion of single particles to folding of proteins that are more complex with the multidimensional folding landscape. Although Kramers’ theory was originally developed for barrier crossing in a one-dimensional potential with a single barrier, experiments and simulations suggest the theory holds for dynamic processes in biomolecules. Interestingly, following the theoretical study, establishing that folding rates (*k*_F_s) of proteins vary as *k*_F_ ~ 1/*η*,^14^ experiments on cold shock protein,^18^ chymotrypsin inhibitor,^19^ and protein L^20^ confirmed Kramers’ high-*η* predictions. Although these studies showed that the rate dependence on *η* follows Kramers’ prediction, this was most vividly demonstrated in single molecule studies only recently by Chung and Eaton.^6^ The success of the Kramers’ theory, which views the complex process of polypeptide chain organization as diffusion in an effective one dimensional landscape, is surprising. However, it has been shown using lattice models^21^ that diffusion in an energy landscape as a function of a collective coordinates, such as the fraction of native contact (*Q*), provides an accurate description of the folding rates obtained in simulations. Subsequently, computational studies^22^ using Gō model for a helix bundle further showed that the rate dependence follows the theoretical predictions including the Kramers turnover, providing additional justification that *Q* is a good reaction coordinate for protein sequences that are well optimized.

In contrast to several studies probing viscosity effects on protein folding and dynamics, frictional effects on nucleic acid folding have been much less studied. Viscogens not only alter the folding rates but also has an effect on the free energies of stabilities between the folding rates. Ansari and Kuznetsov showed that, when corrected for stability changes, the rates of hairpin formation of a DNA sequence was found to be ∝ 1/*η*.^23^ Kramers’ predictions at high *η* were also borne out in the folding of G-quadruplex DNA,^24^ and most recently in the tetraloop-receptor formation in RNA.^25^ These studies show that nucleic acid folding might also be viewed as diffusion in an effective one-dimensional folding landscape.

In this paper, we consider frictional effects on the RNA folding using coarse-grained (CG) simulations. We investigate the variations in rates of folding of a *human* Telomerase hairpin (hTR HP) and an H-type pseudoknot from *Beet western yellow virus* (BWYV PK) as a function of *η*. Because both the HP and PK fold by parallel pathways, our study allows us to examine whether frictional effects affect the flux through parallel pathways in RNA folding. Despite the differences in sequences and the folded structures, the dependence of *k*_F_ on *η* are quantitatively fit using Kramers’ theory including the predicted turnover. The excellent agreement between theory and simulations allows us to estimate a speed limit for RNA, which we find to be (1-4)*μ*s. Surprisingly, we find that the flux through the pathways may be altered by changing solvent viscosity for both the HP and PK. The change in the flux is more pronounced for HP, especially at a temperature below the melting temperature. We argue that this prediction is amenable to experimental tests in RNA even though it has been difficult to demonstrate it for protein folding.

## Materials and Methods

### RNA molecules

We choose a sequence that forms a hairpin (HP) with no tertiary interactions from the human Telomerase (hTR) and a H-type BWYV pseudoknot (PK), which is a minimal RNA motif with tertiary interactions. The folding mechanisms of PKs are diverse,^25^ and they often reach the native structure by parallel pathways.^26^ The use of two RNA molecules with different folded states, with both HP and PK folding occurring by parallel pathways, allows us to examine many consequences of viscosity effects on their folding. The structure of hTR HP (PDB ID 1NA2) has been determined using NMR (see Figure 1A).^27^ The folded structure of the BWYV PK is taken from the crystal structure (PDB ID 437D).^28^ The PK has 28 nucleotides forming two stems. The two loop regions have hydrogen bonding interaction with the stems (Figure 1B). We added an additional guanosine monophosphate to both the 5′ and 3′ terminus to minimize end effects. Thus, the simulated PK has 30 nucleotides.

**Figure 1.**
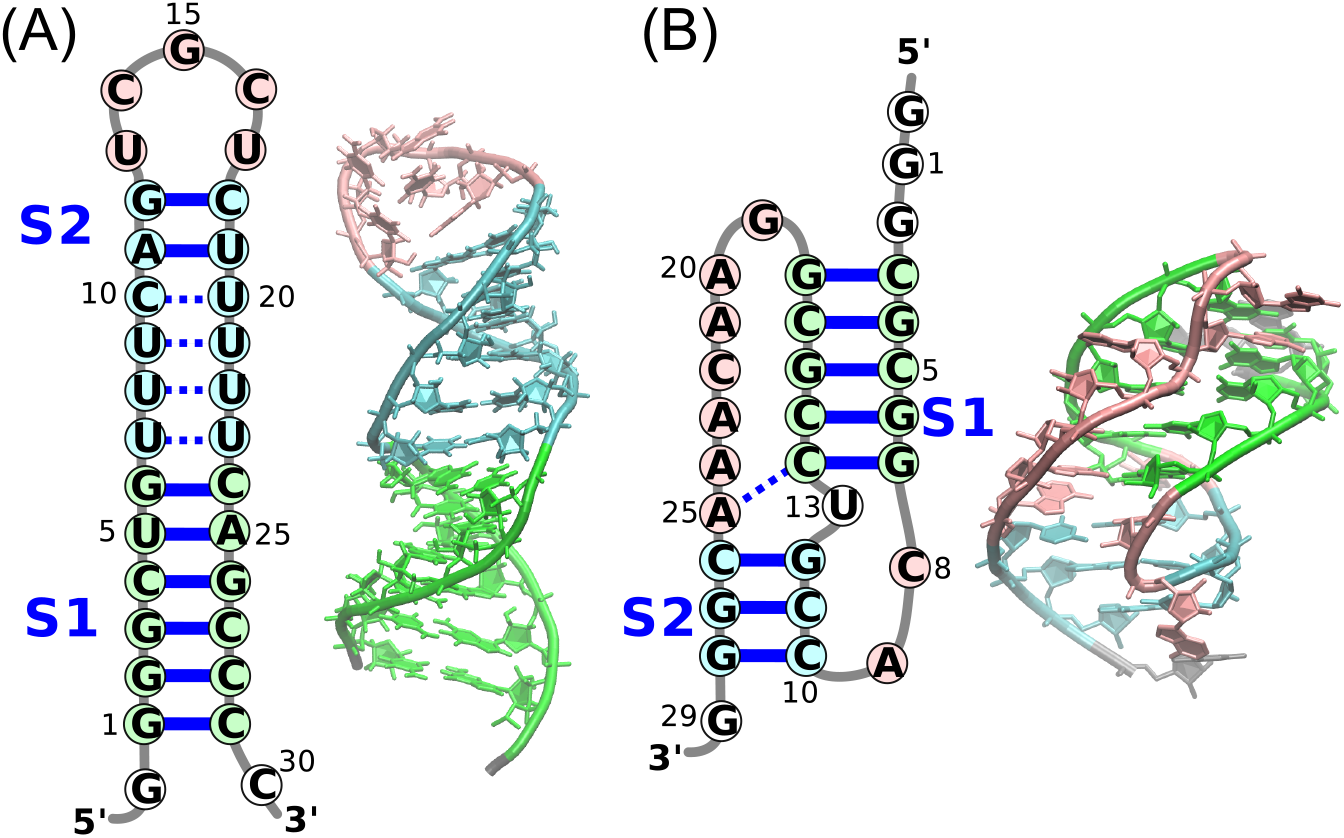
(A) Secondary representation and sequence of human Telomerase hairpin (hTR HP). The folded hairpin structure is on the right. Note that the base pairings in S1 are canonical whereas there are several non-canonical base pairs in S2. (B) Secondary structure of Beet Western Yellow Virus pseudoknot (BWYV PK). The tertiary structure of the PK is shown on the right. In the secondary structures, blue lines represent canonical base pairs (thick lines) and non-canonical pairs (dotted lines).

### Three Interaction Site (TIS) model for RNA

We employed a variant of the TIS model, which has been previously used to make several quantitative predictions for RNA molecules ranging from hairpins to ribozymes.^25,29–31^ We incorporated the consequences of counter ion condensation into the TIS model, allowing us to predict the thermodynamic properties of RNA hairpins and PK that are in remarkable agreement with experiments.^32^ Because the details of the model have been reported previously, we only provide a brief description here. In the TIS model,^29^ each nucleotide is represented by three coarse-grained spherical beads corresponding to phosphate (P), ribose sugar (S), and a base (B). Briefly, the effective potential energy (for details see Ref.^32^) of a given RNA conformation is *U*_TIS_ = *U*_L_ + *U*_EV_ + *U*_ST_ + *U*_HB_ + *U*_EL_, where *U*_L_ accounts for chain connectivity and angular rotation of the polynucleic acids, *U*_EV_ accounts for excluded volume interactions of each chemical group, *U*_ST_ and *U*_HB_ are the base-stacking and hydrogen-bond interactions, respectively.

Electrostatic interactions between the phosphate (P) groups are given by *U*_EL_. The repulsive electrostatic interactions between the P sites are taken into account through the Debye-Hückel theory, 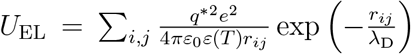, where the Debye length is 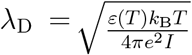. In the present simulations, salt concentration (monovalent ions) is set to 200 mM, which is close to the physiological value. The ionic strength 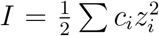 where *c_i_* is the molar concentration, and *z_i_* is the charge number of ion *i*, and the sum is taken over all ion types. Following our earlier study,^32^ we used an experimentally fit function for the temperature-dependent dielectric constant *ε*(*T*).^33^ To account for counter-ion condensation, we used a renormalized charge on the phosphate group, – *q*e*(*q*\* < 1). The renormalized value of the charge on the P group, is approximately given by 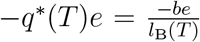, where the Bjerrum length is 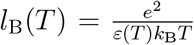, and *b* is the mean distance between the charges on the phosphate groups.^34^ We showed elsewhere^32^ that a constant value of *b* = 0.44 nm accounts for the thermodynamics of several RNA molecules, and is the value adopted here. All the force-field parameters used here are the same as in our earlier study.^32^

### Simulation details

We performed Langevin dynamics simulations by solving the equation of motion,

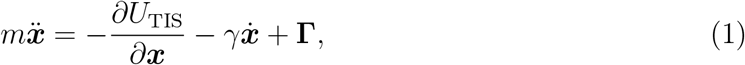

where *m* is the mass of the particle, ***x*** is the coordinate, and **Γ** is a Gaussian random force that satisfies the fluctuation-dissipation relation given by 〈**Γ**_*i*_(*t*)**Γ**_*j*_(*t*′)〉 = 6*γk*_B_*Tδ*(*t* − *t*′)*δ_ij_*. The friction coefficient follows the Stokes-Einstein relation, *γ* = 6*πηR*, where *R* is the appropriate size of the coarse-grained bead (P, S and B) and *η* is the solvent viscosity. The numerical integration is performed using the velocity-Verlet algorithm.^35^

In the high friction regime where *η* = 10^−3^ ~ 10^−2^, we performed Brownian dynamics simulations^36^ by neglecting the inertial term, since the dynamics is over-damped. In this limit, the equation of motion is,

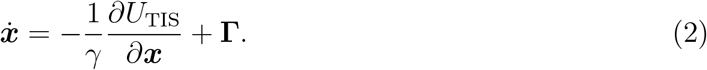

We used reduced units in the analysis of data.^14^ In the TIS representation, we chose the mass of a bead *m* = 116 g/mol, the typical length scale *a* = 0.4 nm, and the energy scale *ε* =1 kcal/mol. Thus, the natural measure for time of Eq. 1 is *τ* = (*ma*^2^)^1/2^ ~ 2ps. In the over-damped condition (Eq. 2), the natural unit of time is 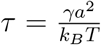. We used this measure to obtain *τ* ≈ 300 ps for converting the simulation times to real times at the viscosity of water, *η_w_* = 10^−3^ Pa · s.^37^

We confirmed that both Langevin dynamics and Brownian dynamics simulations give identical results in the crossover value of *η* = 10^−3^ Pa·s, using simulations of hTR hairpin. The difference between the two simulations method is in the range of statistical error estimated by the jack-knife method. For example, the folding rate for hTR HP is *k*_F_ = 5.5 ± 0.5 ms^−1^ calculated from 100 trajectories generated using Brownian dynamics simulations, and is *k*_F_ = 6.5 ± 0.6 ms^−1^ obtained from another set of 100 trajectories obtained using Langevin dynamics simulations at *η* = 10^−3^ Pa·s.

### Hydrodynamic Interactions

In order to ensure that the results are robust, we did limited simulations of folding by including hydrodynamic interactions (HI). To take into account the effects of HI, we performed Brownian dynamics simulations using the following form with conformation-dependent mobility tensor,

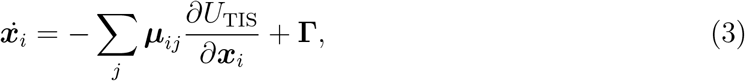

where ***μ***, the mobility tensor, is computed using the Rotne-Prager-Yamakawa approximation,^36^

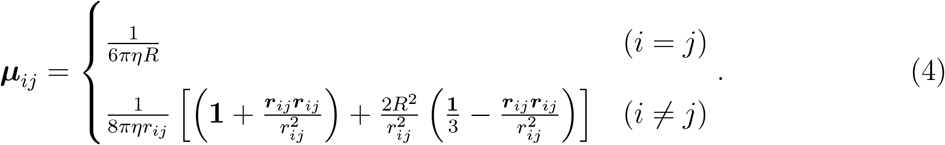

In the above equation, ***r***_*ij*_ is a coordinate vector between beads *i* and *j*. Because coarsegrained beads in our TIS model have different radii (*R*) depending on the type of beads (phosphate, sugar, and bases),^32^ we employed a modified form of ***μ*** developed by Zuk *et al*.^38^

### Thermodynamics properties

We performed temperature-replica-exchange simulations (T-REMD)^39^ to calculate the heat capacity. Temperature was distributed from 0 to 120 °C with 16 replicas at 200 mM of salt concentration. The T-REMD simulation is performed using a low friction (*η* = 10^−5^ Pa·s) to enhance the efficiency of conformational sampling.^35^

### Order parameter

In order to determine if a folding reaction is completed, we used the structural overlap function^40^

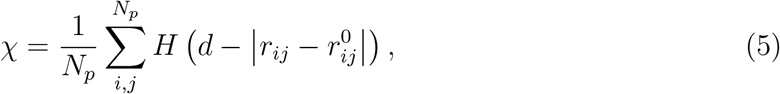

where *H* is the Heaviside step function, *d* = 0.25 nm is the tolerance, and 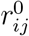 is the distance between particles *i* and *j* in the native structure. The summation is taken over all pairs of coarse-grained sites separated by two or more covalent bonds, and *N_p_* is the number of such pairs. The structural overlap function quantifies the similarity of a given conformations to the native conformation. It is unity if the conformation is identical to the native state. In *T*-quench kinetics simulations, if the value of the structural overlap function exceeds a threshold, 〈*χ*〉_*T*_L__, the trajectory is deemed to be completed, and the folding time *τ_i_* is recorded; 〈*χ*〉_*T*_L__ is the thermodynamic average at the lower simulation temperature at which RNA molecules are predominantly folded (Table 1). In addition to *χ*, we also calculated the average value of 〈*R*_g_〉, measurable in scattering experiments (SAXS or SANS), to assess the temperature dependence of compaction of the RNA molecules.

**Table 1:**
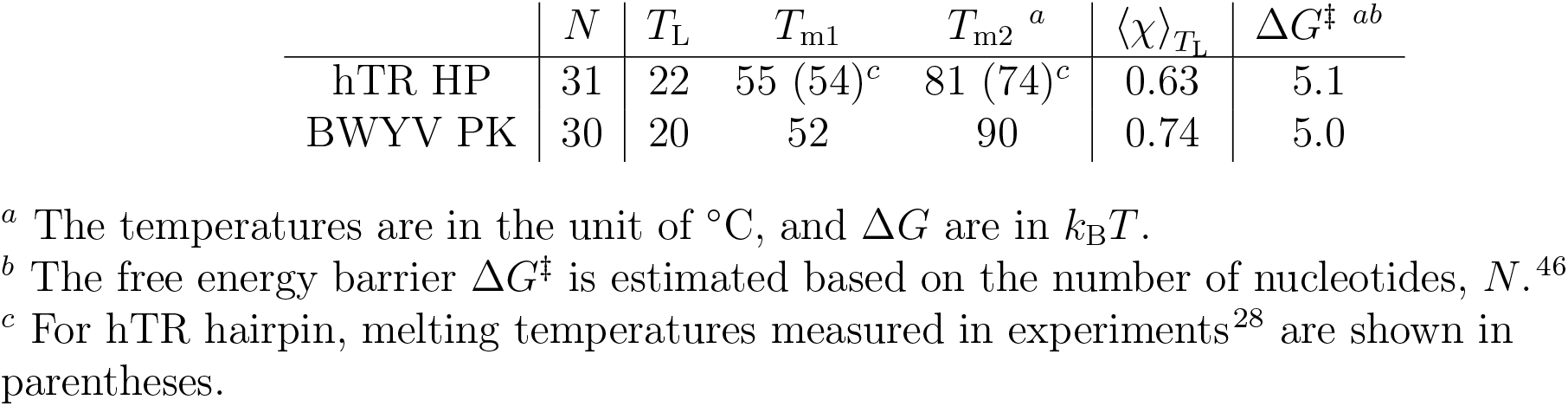
Thermodynamic Properties of the RNAs.

| | $N$ | $T_L$ | $T_{m1}$ | $T_{m2}^a$ | $\langle\chi\rangle_{T_L}$ | $\Delta G^\ddagger^{ab}$ |
| --- | --- | --- | --- | --- | --- | --- |
| hTR HP | 31 | 22 | 55 (54)<sup>c</sup> | 81 (74)<sup>c</sup> | 0.63 | 5.1 |
| BWYV PK | 30 | 20 | 52 | 90 | 0.74 | 5.0 |

<sup>a</sup> The temperatures are in the unit of $^\circ\text{C}$ , and $\Delta G$ are in $k_B T$ .
<sup>b</sup> The free energy barrier $\Delta G^\ddagger$ is estimated based on the number of nucleotides, $N$ .<sup>45</sup>
<sup>c</sup> For hTR hairpin, melting temperatures measured in experiments<sup>27</sup> are shown in parentheses.

### *T*-quench folding

To prepare the initial structural ensemble for *T*-quench simulations, we first performed low-friction Langevin dynamics simulations. The simulation temperatures are chosen to be 1.2 times higher than the second melting temperature (in Kelvin unit) to ensure that completely unfolded conformations are populated. After generating a sufficiently long trajectory to ensure that the chain has equilibrated, the unfolded conformations are sampled every 10^5^ time steps. Finally, we collected hundreds of conformations which were used as initial structures in the *T*-quench folding simulations.

In order to initiate folding, starting from an unfolded structure, we quench the temperature to *T*_S_, and generated folding trajectories using Langevin or Brownian dynamics simulations by varying the solvent viscosity from *η* = 3.2 × 10^−9^ to 10^−2^ Pa·s (cf. water viscosity *η*_w_ ~ 10^−3^ Pa·s). The viscosity is directly related to the friction coefficient as *γ* = 6*πηR* where *R* is the radius of coarse-grained beads. For each condition, at least 100 folding trajectories are generated. Folding time *τ_i_* is measured by monitoring the overlap function, *χ*, in each trajectory *i*, and folding rates were calculated by averaging over *M* trajectories, 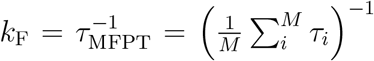.^42^ We used two values of *T*_S_. One is *T*_S_ = *T*_m1_, which is the lower melting temperature in the heat capacity curve (Figure 3), and the other is *T*_S_ = *T*_L_ < *T*_m1_. The values of *T*_L_ and *T*_m1_ are listed in Table 1.

### Data analysis using Kramers’ rate theory

The simulation data for RNA is analyzed using Kramers’ theory^16^ in which RNA folding is pictured as a barrier crossing event in an effective one-dimensional landscape (see Figure 2). For our purposes here it is not relevant to determine the reaction coordinate because we estimate the values of the barrier heights theoretically.^42^ According to transition state theory (TST), the reaction rate is expressed as

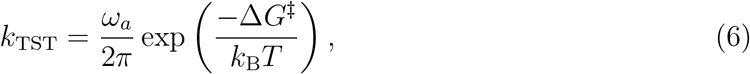

where *k*_B_ is the Boltzmann constant, and *T* is the temperature. The rate, *k*_TST_, gives us an upper bound of the true reaction rate since, in the TST, there are no recrossing events once RNA reaches the saddle point.^1,2^

**Figure 2.**
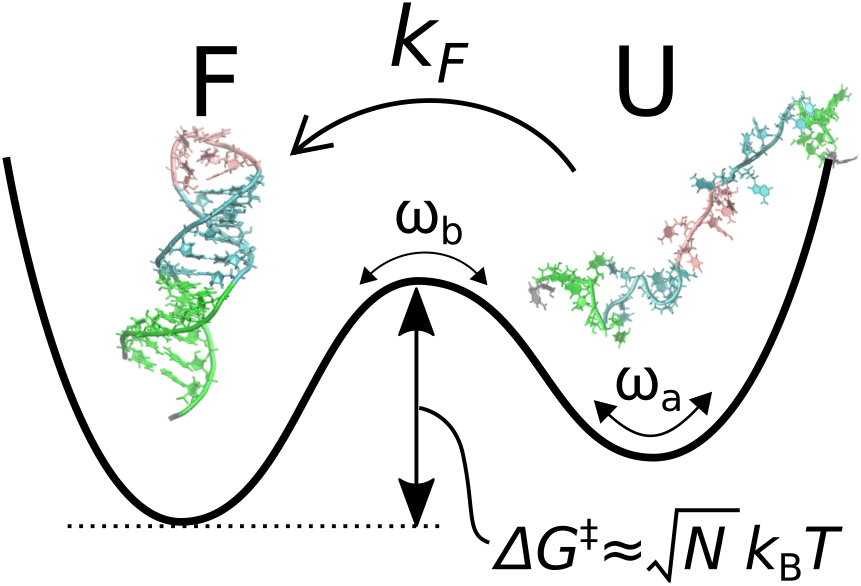
Schematic of effective one-dimensional landscape of RNA folding from the unfolded state (U) to the folded state (F). Parameters, *ω_a_* and *ω_b_*, determine curvatures of the free energy surface at the reactant basin (U) and the saddle point. Δ*G*^‡^ is the height of the free energy barrier.

The Kramers’ folding rate can be written as,

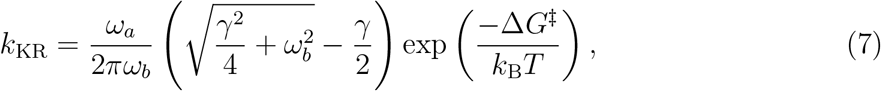

where *γ* is the friction coefficient, *ω_a_* and *ω_b_* are the parameters that determine curvatures of the free energy surface at the reactant basin and saddle point (maximum in the free energy surface), respectively (Figure 2). It is assumed that in the vicinity of the saddle point, the free energy may be approximated by a parabola 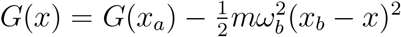 where *x* is an unknown reaction coordinate, *x_b_* is the position of the saddle, and 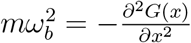.

In the high friction limit,

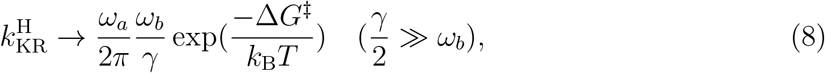

which shows that the folding rate should depend on the inverse of the friction coefficient. When the friction is small, the rate linearly approaches the rate determined by the TST, 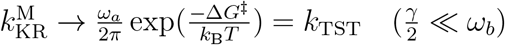.

If we further consider the time scale at which local equilibrium is achieved (very weak damping limit), the rate becomes

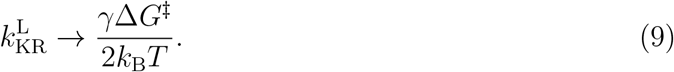

In this regime, barrier crossing is controlled by energy diffusion,^44^ and the TST is no longer valid. These extremely well known results, used to analyze the simulations, can be summarized as follows: Kramers’ theory predicts that 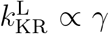 in the low friction regime, *k*_KR_ reaches a maximum in moderate friction 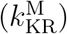, and 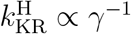 in the high friction regime.

### Barrier heights in the folding landscape

In order to use Eq. 7 to analyze simulation data the free energy barrier to folding has to be calculated. However, estimating barrier heights is non-trivial in complex systems because the precise reaction coordinates is difficult, particularly for RNA in which ion effects play a critical role in the folding reaction. In order to avoid choosing a specific reaction coordinate, we appeal to theory to calculate the effective barrier height. One of us has shown,^42^ which has been confirmed by other studies,^44^ that for proteins the free energy barrier ≈ 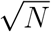 where *N* is the number of amino acids. In the context of RNA folding, we showed that there is a robust relationship between the number of nucleotides (*N*) and the folding rates, *k*_F_ ≈ *k*_o_exp(−*αN*^0.5^).^45^ Experimental data of folding rates spanning seven orders of magnitudes (with *N* varying from 8 to 414) were well fit to the theory using *α* = 0.91 and 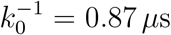 (speed limit for RNA folding).^45^ Therefore, in this study, we estimate the free energy barrier based on *N* alone. A clear advantage of using the theoretical estimate is that it eliminates the need to devise a reaction coordinate. The values of the barrier height for the two RNAs are given in Table 1.

## Results

### Thermal denaturation

We calculated the heat capacities of the HP and the PK at the monovalent salt concentration of 200 mM (Figure 3). The heat capacities have two distinct peaks, which indicate there is at least one distinct intermediate between the unfolded and folded states. This finding is consistent with previous experimental and simulation studies.^27,46,47^ From the position of the peaks, we determined the two melting temperatures, *T*_m1_ and *T*_m2_, whose values are listed in Table 1. It should not go unnoticed that the melting temperatures, *T*_m1_ and *T*_m2_, for the hTR HP are in excellent agreement with experiments, demonstrating the effectiveness of TIS model in predicting the thermodynamic properties of RNA. The experimental heat capacity curve is not available for BWYV PK at 200 mM salt concentration, and hence the values reported in Table 1 serve as predictions.

**Figure 3.**
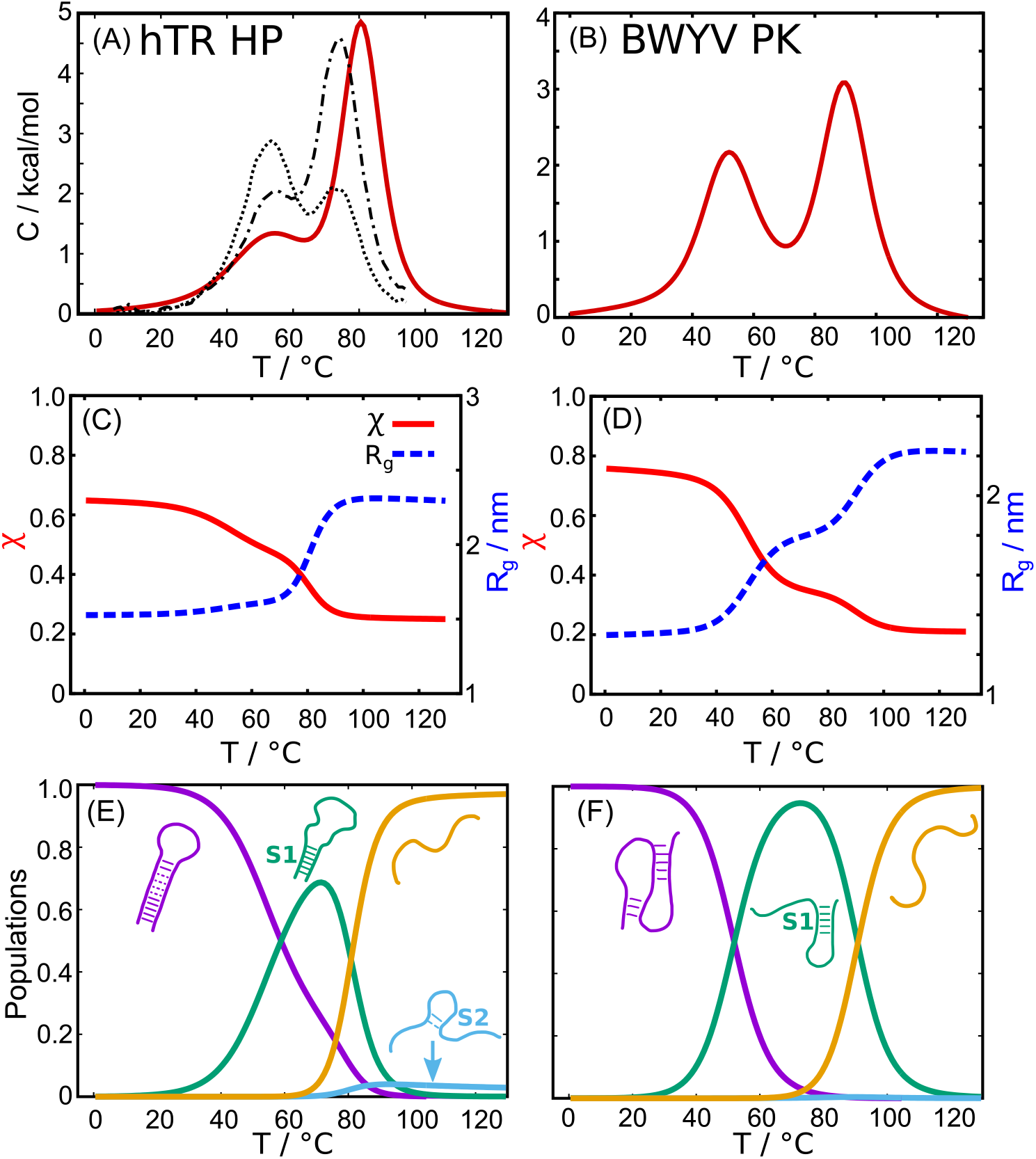
Temperature dependence of thermodynamic properties at 200 mM of monovalent salt concentration. (**A, B**) Heat capacity of (A) hTR HP and (B) BWYV PK. The red lines are heat capacities, *C*(*T*), computed from the T-REMD simulations. Black lines in (A) are UV absorbance melting profiles (*δA/δT*) at 260 nm (dotted line) and 280 nm (dot-dash line) experimentally reported elsewhere.28 The scale of *δA/δτ* is not relevant because we only compare the positions of the peaks. The melting temperatures for the HP are given in Table 1. For the PK, the values of *T_m_*s are predictions. (**C, D**) Structural overlap functions (*χ*; red solid) and radius of gyration (*R_g_*; blue dashed). (**E, F**) Populations of folded (purple), intermediate (green and cyan), and unfolded states (yellow) as functions of *T*.

### Viscosity dependence of the folding rates, Kramers turnover and absence of internal friction

Friction dependent folding rates obtained from the *T*-quench simulations are shown in Figure 4. In the high friction regime, *η* ≳ 10^−5^ Pa·s, the folding rates *k*_F_ decrease as the friction is increased. This behavior is found in both HP and PK at both temperatures, *T*_L_ and *T*_m1_. In the moderate friction regime, 10^−7^ ≲ *η* ≲ 10^5^ Pa·s, the folding rates reach maximum values. For *η* ≥ 10^−6^ Pa·s, we fit the values of *k*_F_ to Eq. 7 with 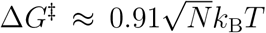. By adjusting the two free parameters, *ω_a_* and *ω_b_*, Eq. (7) quantitatively accounts for the simulation data (lines in blue in Figure 4, parameters are summarized in Table 2). Thus, the variation of *k*_F_ ∝ *η*^−1^ in the high friction regime shows that Kramers’ theory accurately describes the dependence of the folding rates on *η* of these two RNA constructs. We conclude that even for RNA, driven by electrostatic interactions, folding could be pictured as diffusive process in an effective one dimensional landscape. As *η* decreases there is a maximum in the rate followed by a decrease in *k*_F_ at low *η*, which shows the expected Kramers turnover. The quantitative account of simulation data on *k*_F_ using Kramers’ theory at high *η* shows absence of internal friction in the folding process of these RNA constructs.

**Table 2:**
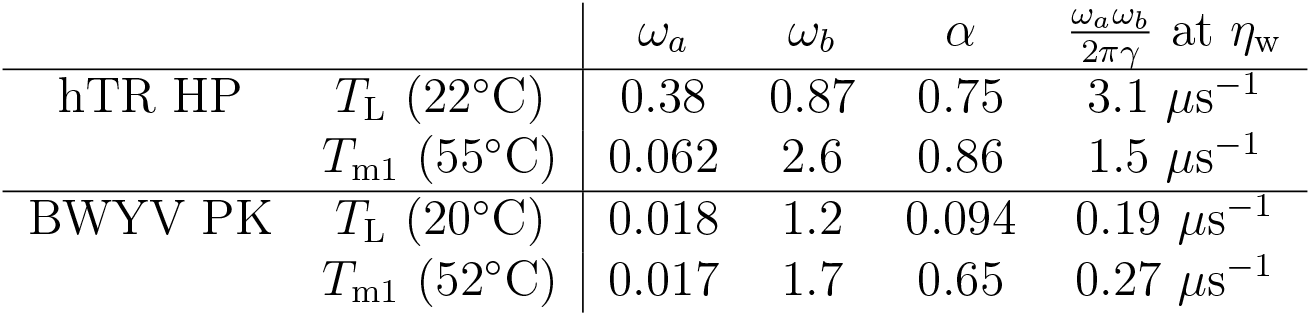
Fitting Parameters.

| | | $\omega_a$ | $\omega_b$ | $\alpha$ | $\frac{\omega_a \omega_b}{2\pi\gamma}$ at $\eta_w$ |
| --- | --- | --- | --- | --- | --- |
| hTR HP | $T_L$ (22$^\circ\text{C}$) | 0.38 | 0.87 | 0.75 | 3.1 $\mu\text{s}^{-1}$ |
| | $T_{m1}$ (55$^\circ\text{C}$) | 0.062 | 2.6 | 0.86 | 1.5 $\mu\text{s}^{-1}$ |
| BWYV PK | $T_L$ (20$^\circ\text{C}$) | 0.018 | 1.2 | 0.094 | 0.19 $\mu\text{s}^{-1}$ |
| | $T_{m1}$ (52$^\circ\text{C}$) | 0.017 | 1.7 | 0.65 | 0.27 $\mu\text{s}^{-1}$ |

In the low friction regime, the dependence of *k*_F_ ∝ *η^α^* with a positive *α* (lines in purple in Figure 4, and values of *α* are in Table 2). In contrast to the high friction case, the low *η* dependence varies with each RNA molecule, and deviates from the expectations based on Kramers theory, which predicts *α* ≈ 1.

**Figure 4.**
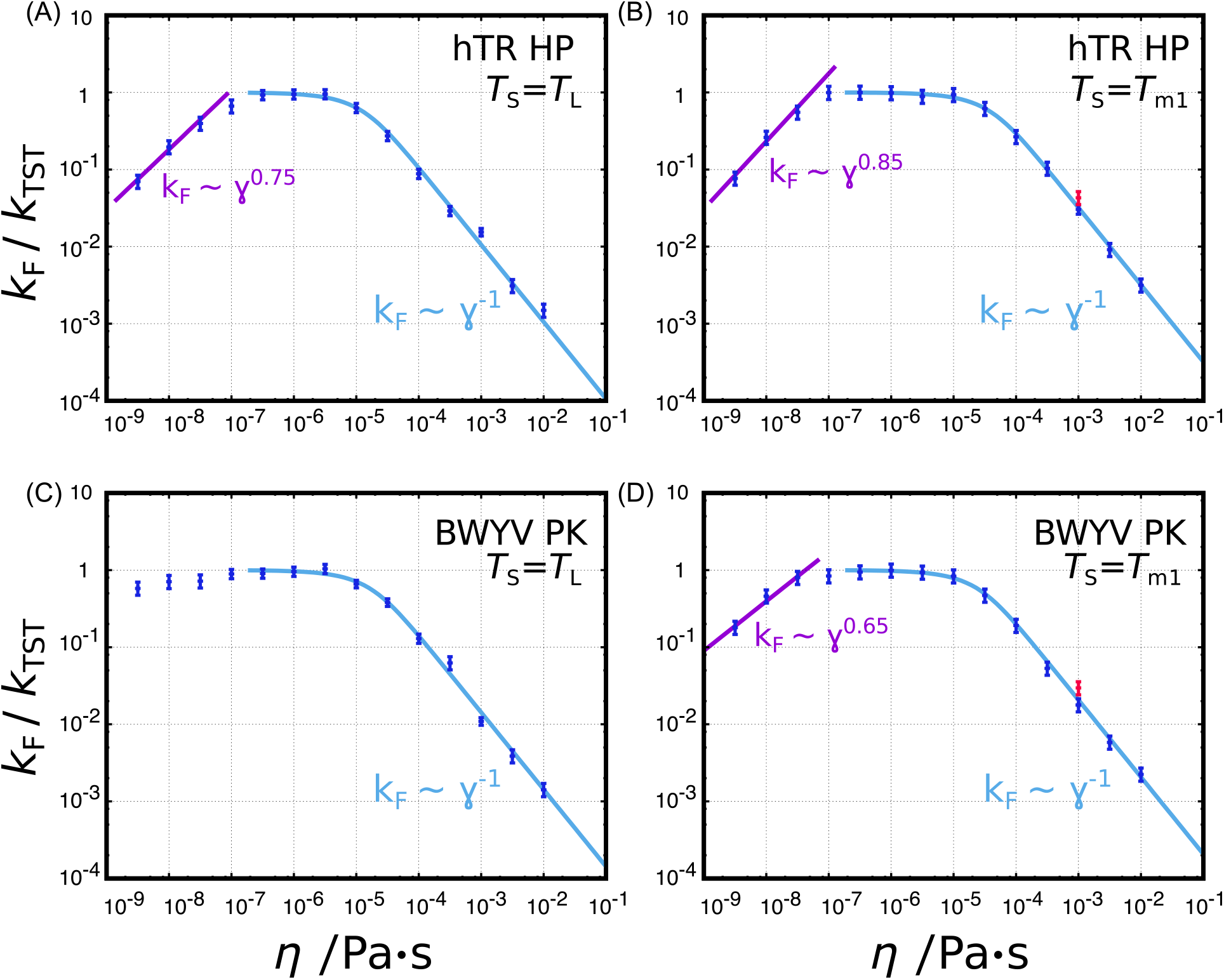
Friction (viscosity) dependence of folding rates at simulation temperatures *T*_S_ = *T*_L_ and *T*_m1_ for hTR HP (A and B) and BWYV PK (C and D). See Table 1 for the numerical values of *T*s. Folding rates (blue bars) are normalized by the values from the transition state theory, *k*_TST_ (Eq. 6). The error bars indicate 95% confidence level. The data in the moderate to high friction regime (*η* ≥ 10^−6^ Pa·s) were fit to Eq. 7 (lines in light blue). The data in the low friction regime (*η* ≤ 10^−7.5^ Pa · s) were fit to Eq. 9 except rates of BWYV PK at *T*_L_. The results of Brownian dynamics simulations with hydrodynamic interactions at the water viscosity (*η* = 10^−3^ Pa·s) are shown in red in (B) and (D) for hTR HP and BWYV PK, respectively.

### Viscosity effects on hairpin folding pathways

The hTR HP has two regions of consecutive canonical base pairs, which we label Stem 1 (S1) and Stem 2 (S2) (Figure 1A). Four non-canonical base pairs are flanked by S1 and S2. Because of the differences in base pairing between these regions, the folding pathways may be visualized in terms of formation of S1 and S2 separately. It is clear that S1 is more stable than S2, and we expect the former to form first in the folding process according to the stability hypothesis suggested by Cho, Pincus, and Thirumalai (CPT).^25^ In order to assess if the difference in stability leads to friction-induced changes in the flux between the two pathways (S1 forms before S2 or vice *versa*), we calculated the fraction of pathways (Φ) from the folding trajectories which is obtained by counting the number of trajectories that reaches the folding by first forming S1. We found different trends between the two temperatures (Figure 5). At *T*_m1_, the dominant pathway (labeled I) is characterized by formation of the more stable S1 at all values of *η*. The flux through I is ≈ 0.8 at *η* values close to *η*_w_ (the water viscosity) and minor pathway (II) (1 − Φ) ≈ 0.2 in which S2 forms first followed by S1 (figure 5A). This finding is in accord with the expectation based on the relative stabilities of S1 and S2.^26^ The dominance of pathway I at *T*_m1_ suggests that folding starts away from the loop with the formation of base pair between nucleotides G1 and C29 and the HP forms by a zipping process.

**Figure 5.**
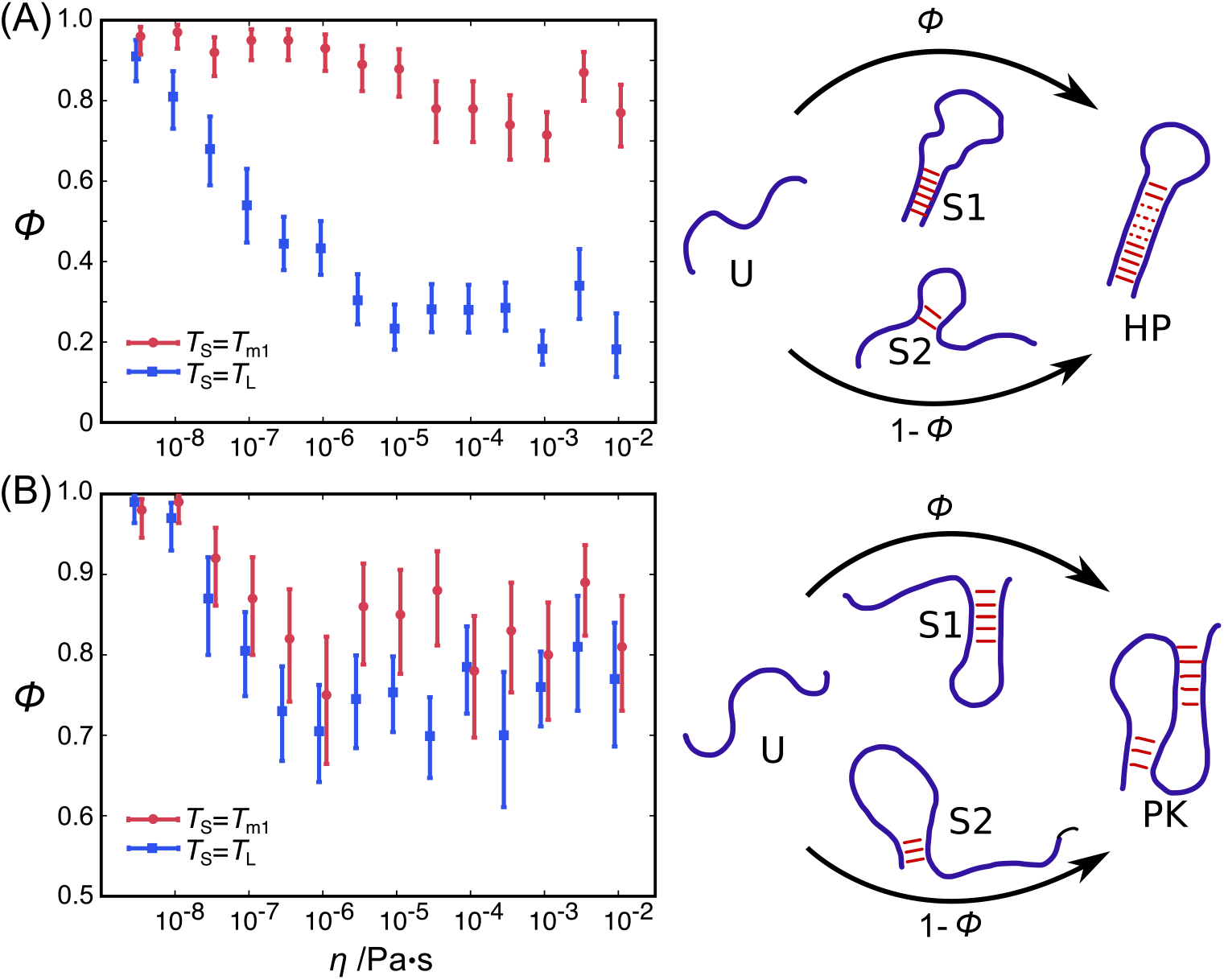
Variations in the flux through the two pathways as a function of viscosity. (A) hTR HP and (B) BWYV PK. The definition of states for each RNA is schematically shown on the right. (**A**) hTR HP structure naturally splits into two helices, S1 and S2 (Figure 1A). The folding pathways are classified either if S1 forms first or S2 forms first. The fraction Φ is the number of trajectories in which S1 forms first divided by the total number of trajectories. (B) BWYV PK has two hairpin stems, S1 and S2, allowing us to classify the pathways in the same manner as in A. The error bars indicate 95% confidence intervals.

Interestingly, at the lower temperature *T*_L_, we find that Φ changes substantially as *η* increases (Figure 5). At *η* in the neighborhood of *η*_w_, Φ is only ≈ 0.2, which implies that at *T*_L_ folding predominantly occurs in the less dominant pathway (II), by first forming the less stable S2. This finding may be understood using our previous study on P5GA, a 22-nucleotide RNA hairpin containing only WC basepairs.^38^ We found that, although there are multiple ways for P5GA to fold, the most probable route is through formation of a short loop (SL) that initiates nucleation of base pair formation involving nucleotides close to the loop. With that finding in mind, we can rationalize the flux changes at *T*_L_. The entropy loss (Δ*S*) due to loop closure, which in hTR HP would being the two Uracil bases (Figure 1) close enough to initiate G-C base pair (nucleation step), would be small (*T*Δ*S*≈*k*_B_*T*ln5). Once the G-C base pair near the loop forms, zipping occurs leading to HP formation. At *T*_m1_ > *T*_L_, S1 formation occurs first which necessarily involves long loop (LL) formation that brings 5′ and 3′ ends close. At high temperature this process is facile even though *T*Δ*S* ≈ *k*_B_*T* ln30. When the 5′ and 3′ are close, the highly favorable enthalpy gain due to the formation of a number of favorable WC basepairs compensates for the entropy loss due LL formation.

The argument given above to explain the Φ values at *T*_L_ can be substantiated by analyzing a typical folding trajectory at *η*_w_ = 10^−3^ Pa · s shown in Figure 6. Before folding occurs, there are several (three times in this particular trajectory) attempts to form S2 involving the favorable SL, as found in P5GA hairpin.^37^ This step is the expected initiation step in helix nucleation. However, formation of S2, needed for growth of the helix, is disrupted because S2 is inherently unstable. Consequently, I2 unfolds and pauses in that state for a long time (Figure 6). In the fourth attempt, the formation of two base pairs near the loop is followed by formation of the non-canonical base pairs, followed by S1, resulting in the folding of the HP. Interestingly, the transient S2 formation is only observed at higher friction regime (Figure 6B inset). At *η* > 10^−4^ Pa · s, there are, on average, 5 ~ 10 attempts of S2 formation before the RNA folds, whereas it does not apparently occur at lower *η*, which is dominated by energy diffusion.

**Figure 6.**
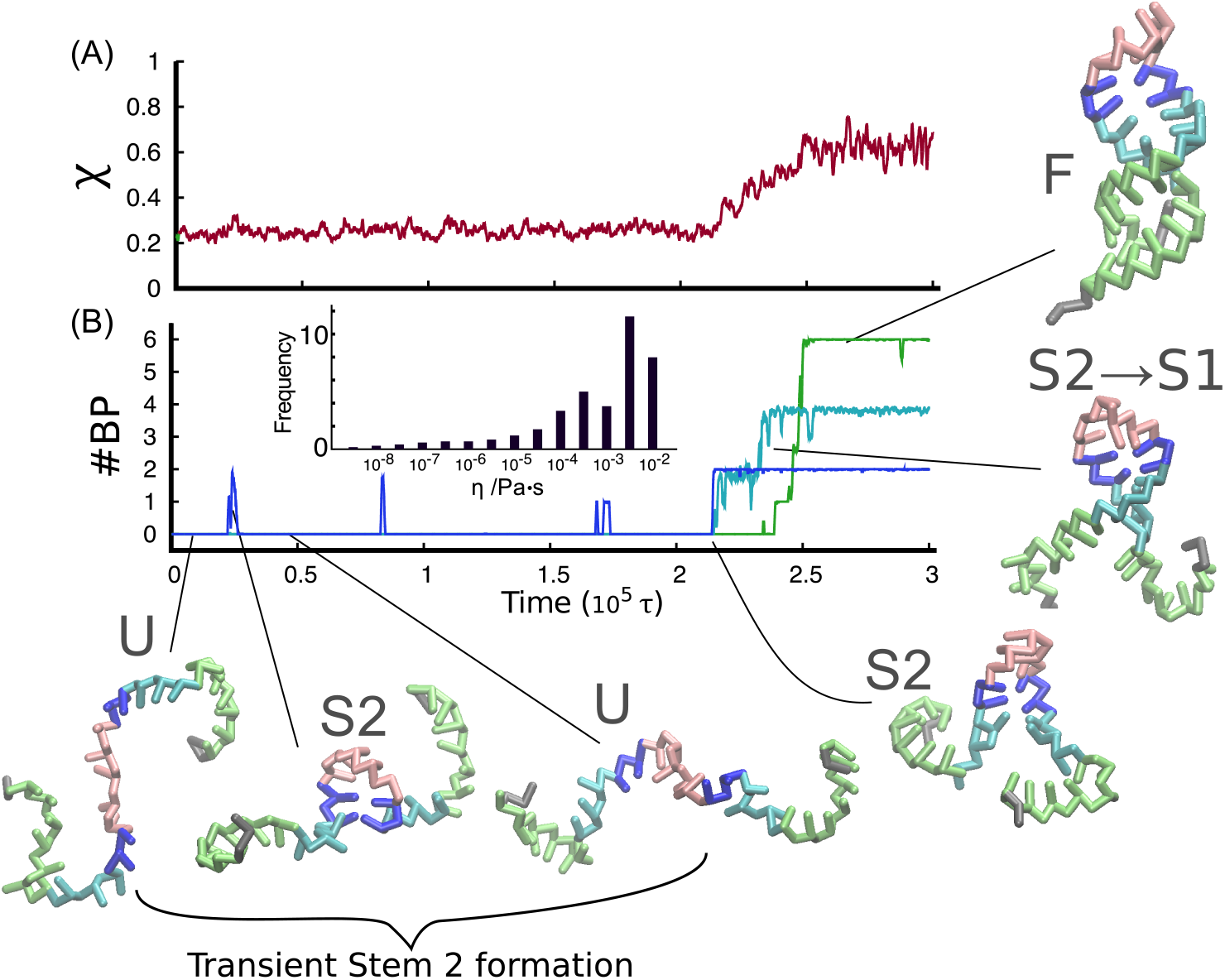
A typical folding trajectory of hTR HP simulated at high viscosity (*η* = 10^−3^ Pa · s) with *T*_S_ = *T*_L_. Time series in (A) shows the structural overlap function and (B) number of basepairs formed in each stem regions (blue: Stem 2, cyan: Non-canonical, green: Stem 1) are shown along with several snapshots of representative conformations. In this trajectory, S2 transiently forms three times before the RNA folds. The folding was initiated with the formation of S2 and followed by the non-canonical basepairs and S1 at last. (B inset) Averaged number of the transient S2 formations before hTR HP reaches the folded state depending on *η*.

### Viscosity alters the flux through the parallel routes in BWYV PK folding

In BWYV PK folding, there are two potential intermediates, I1 characterized by the formation of the more stable Stem 1, or I2 where only Stem 2 is formed. In Figure 5(B), we show Φ as a function of viscosity. In contrast to the hTR HP case, the pathway through I1 is always dominant at all values of *η* at both the temperatures. At the viscosity of water, the fraction Φ ≈ 0.8. This result is consistent with experimental studies indicating that I1 is the major intermediate.^48,49^ Our previous study also showed that the thermal and mechanical (un)folding occur predominantly through the I1 state.^47^ The present results show that I1 state is not only thermodynamically stable, but also is the major kinetic intermediate. Folding of the PK, which occurs by parallel pathways, with the dominant one being *U* → *I*1 → *F*(Φ ≈ 0.8 at *η*_w_ for example). In contrast to hTR HP, the loop entropy in the PK is comparable (Figure 1), and hence the flux between the two pathways is determined by the CPT stability hypothesis.^25^

In the dominant pathway, the folding occurs by the following two steps (see Supplemental Movie): (i) Stem 1 folds rapidly after *T*-quench (〈*τ*_U→1_〉 = 0.02ms at *η* = 10^−3^) forming the intermediate (I1) state, and then (ii) Stem 2 folds after a substantial waiting time (〈*τ*_I→F_) = 0.95ms). Since there is a large gap in the time scale between the two transitions, the rate of the whole process (U→F) is dominated by the second rate determining phase (*τ*_MFPT_ ≈ 1 ms).

### Frictional effects on individual steps in folding

We have already shown the rates for the whole folding process (U → F) of BWYV PK depends on the viscosity in accord with Kramers’ theory (Figure 4). Since there is a major intermediate, I1, in the reaction process, we analyzed the folding rates by decomposing folding into two consecutive reactions, U → I and I → F. Figure 7 shows the frictional dependence of the folding rates for the two transitions; *k*_I→F_ shows almost the same behavior as *k*_U→F_ since the two time scales are essentially the same (compare Figure 7 and Figure 4 C and D). It is interesting that the rate of the faster transition, *k*_U→I_, also exhibits the Kramers-type dependence especially in the high friction regime, i.e. *k*_F_ ∝ *η*^−1^ for *η* ≳ 10^−5^ Pa·s. This result indicates that, even if the folding reaction involves intermediates, (i) the entire rate still exhibits the Kramers-type dependence, at least in a case that one of the substeps is rate limiting, and (ii) a substep which is not rate determining to the entire rate constant, may also show Kramers-type viscosity dependence.

**Figure 7.**
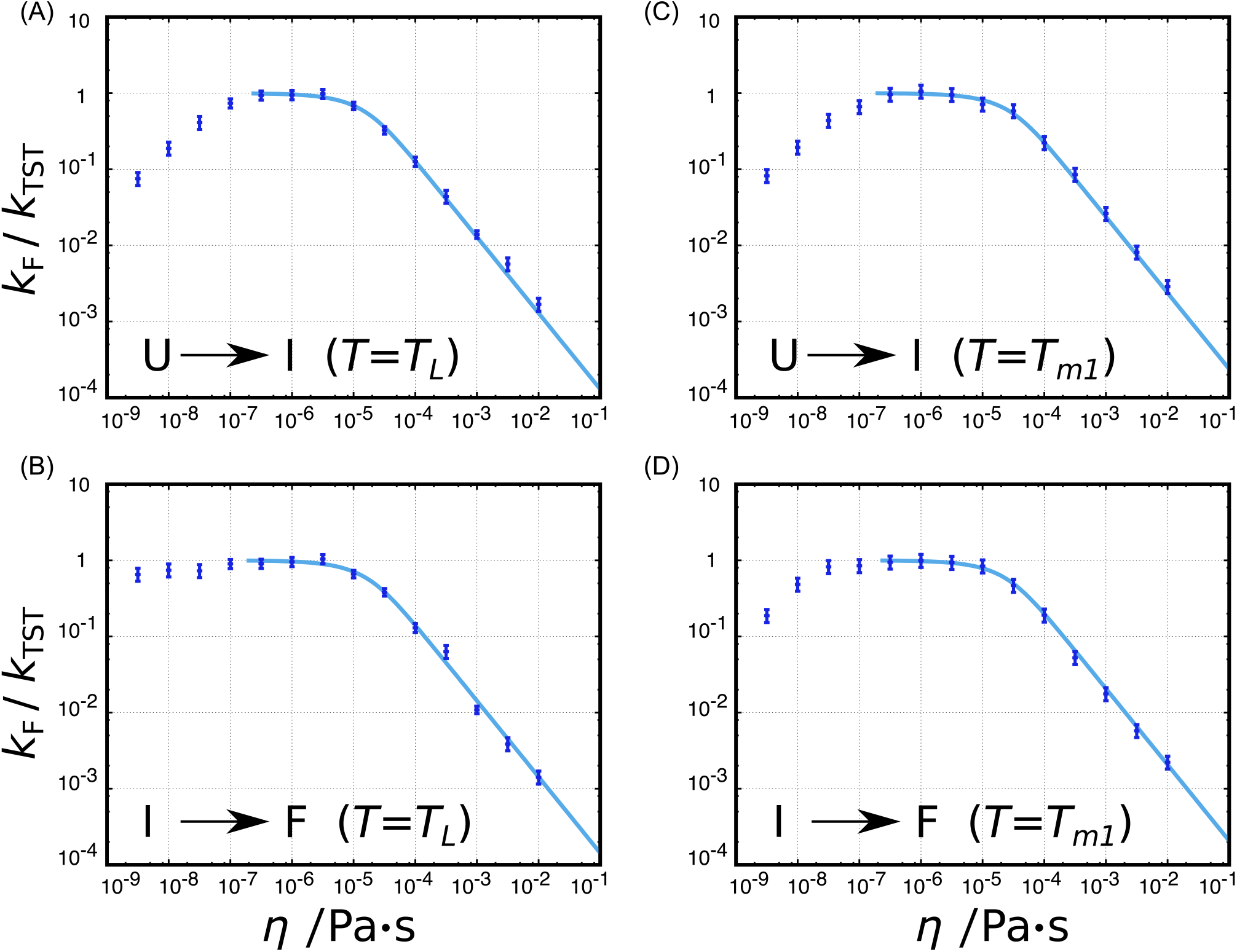
For BWYV PK, folding rates are individually calculated for two sequential structural transition through the intermediate, U → I (middle panels) and I → F (lower panels). The results of the whole process U → F are shown in Figure 4 (C, F).

## Discussion

### Effect of hydrodynamic interactions

In order to ensure that our conclusions are robust, we also examined the effect of hydrodynamic interactions by performing simulations only at the water (*η*_w_ = 10^−3^ Pa·s) for both hTR HP and BWYV PK. As shown in Figure 4 (red bars in B and D), the hydrodynamic interaction (HI) accelerates the folding rates, but its effect is not as significant as changing the viscosity. At *η*_w_, hTR HP folds with *k*_F_ ~ 9.5 ms^−1^ with HI, whereas *k*_F_ ~ 6.5 ms^−1^ without HI. Thus, the reaction is about 1.5 times faster if HI is included. In BWYV PK case, *k*_F_ ~ 1.9 ms^−1^ with HI, whereas *k*_F_ ~ 1.1ms^−1^ without HI, leading to a factor of ~1.7 increase, which is similar to the hTR HP case.

### Changes in Viscosity alter the flux between parallel assembly of RNA

It is well accepted that RNA in general, and PK in particular, fold by parallel pathways.^25,26,50^ Recently, it was shown unambiguously that monovalent cations could change the flux to the folded state between the two pathways in the VPK pseudoknot. Surprisingly, we find here (see Figure 5) that Φ could be also altered by changing the viscosity for both the HP and the PK. Although the same prediction was made in the context of protein folding,^14^ it is difficult to measure *η* dependence of Φ because the secondary structures in proteins are not usually stable in the absence of tertiary interactions. This is not the case in RNA. For instance, S1 and S2 are independently stable and hence their folding could be investigated by excising them from the intact RNA. Consequently, Φ as a function of *η* can be measured. Based on the results in Figure 5 showing that by varying *η* or *η* and *T*, our prediction could be tested either for the hTR HP or the extensively studied PK (BWYV or VPK). For example, at *T*_L_ we find that Φ changes from 0.2 to 0.4 as *η* is varied over a broad range for hTR HP. Although not quite as dramatic, the changes in Φ are large enough for BWYV PK to be detectable. The stabilities of the independently folding S1 and S2 constructs can be also altered by mutations. For instance, by converting some of the non-canonical base pairs neighboring S2 to WC base pairs in the hTR HP would increase the stability of S2. Because there are a variety of ways (concentration of ions, temperature, and mutations) of altering the independently folding units of RNA, our prediction that Φ changes with *η* could be readily tested experimentally.

### Speed limit for RNA folding

Based on the idea that a protein cannot fold any faster than the fastest time in which a contact between residues that has the largest probability of forming, it has been shown that the speed limit (*τ_SL_*) for protein folding is *τ_SL_* ≈ 1 *μ*s.^51^ Using the observation that the typical folding barrier height scales as 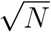 (see Eq. (10) in^43^) and analyses of experimental data^52^ it was shown that *τ_SL_* ≈ *τ*_0_ ≈ (1−10)*μ*s, where 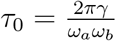 is the inverse of the prefactor in Eq. 8. A similar style of analysis of the experimental data shows that for RNA *τ_SL_* ≈ 1 *μ*s.^45^ Here, an estimate of 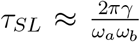 using the values of *ω_a_* and *ω_b_* in Table 2 and *γ* corresponding to water viscosity yields 0.7 *μ*s for the HP and 3.7 *μ*s for the PK. Alternatively, the value of 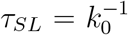 where *k*_0_ = *k*_F_ exp(−0.91*N*^0.5^)(*k*_F_ is the folding rate obtained using simulations) gives *τ_SL_* ≈ 1 *μ*s for the HP and *τ_SL_* ≈ 6 *μ*s for the PK. If *τ_SL_* is equated with the transition path time then we can compare to estimates made for DNA hairpins^53^ and for RNA constructs (several PKs and the *add* riboswitch)^54^ obtained using single molecule experiments. The values range from about (1 ~ 10) *μ*s. Thus, there are compelling reasons to assert from the present and previous theoretical and experimental studies that an RNA cannot fold any faster than about 1 *μ*s.

### Influence of Dielectric Friction

In this article we have treated the electrostatic interactions implicitly, and hence only systematic and viscous dissipative forces act on the interaction sites of RNA. We have not considered the effects of dielectric friction, which could be significant even for an ion moving in an electrolyte solution.^55–58^ In RNA folding, the many body nature of the problem makes it difficult to estimate the magnitude of the dielectric friction. There are multiple ions, with significant ion-ion correlations, that condense onto the RNA in a specific manner dictated by the architecture of the native fold.^31^ The magnitude of dielectric friction in this many body system of highly correlated ion could be significant, which in turn could affect the kinetics of RNA folding. Despite this important issue, which has not been investigated to our knowledge, it is comforting to note that experiments as well as simulations reporting viscosity effects on RNA folding appear to be in accord with Kramers’ theory.

### Transmission Coefficients

The ratio, 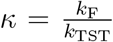 shown in Figure 4, can be as small as ≈ 10^−3^, in the high viscosity region. Recently, based on transition path velocity as a measure of recrossing dynamics^59^ the values of *κ* have been measured in single molecule pulling experiments^60^ for several DNA hairpins. By fixing the mechanical force at the transition midpoint, where the probability of being folded and unfolded are equal, the folding trajectories were used to estimate that *κ* ≈ 10^−5^.^60^ For RNA hairpins it is known that folding times obtained by *T*-quench are larger by at least one order of magnitude relative to times obtained by quenching the force.^29^ Thus, the calculated values of *κ* are not inconsistent with experiments on DNA hairpins under force. It would be most interesting to examine the viscosity dependence of *k*_F_ by maintaining the RNA molecules under tension.

## Conclusions

Using the TIS coarse-grained model, we investigated thermodynamics and folding kinetics of a hairpin and an H-type pseudoknot RNA molecules, focusing on the dependence of the folding rates on the solvent viscosity. From temperature-quench folding simulations, we showed that the folding rates follow the so-called Kramers turnover; the rate increases in the low friction regime and decreases at high friction, with a maximum rate at moderate friction. For both the hairpin and the pseudoknot, the dependence of the folding rates between moderate to high friction regime is robust, and is in accord with the Kramers’ theory. We find clear *η*^−1^ dependence in the folding rates, leaving little doubt that RNA folding involves a diffusive search in an effective low dimensional folding landscape.

A major potentially testable prediction is that in the *η* values that are accessible in experiments the flux between pathways by which RNA folds depends on *η*. Because the stabilities of the individual stems could be altered in RNA easily, our prediction is amenable to experimental test.

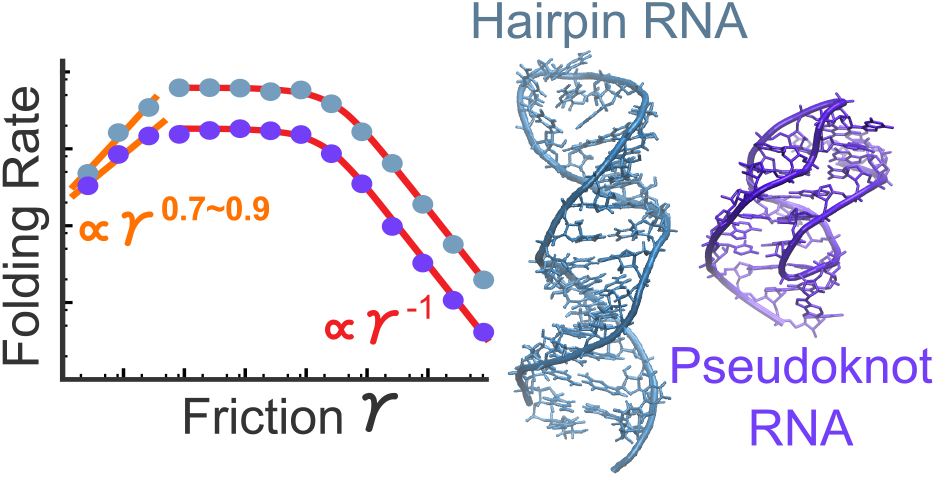

## Acknowledgement

DT would like thank Bill Eaton for numerous discussions over the years about the role of friction in many problems involving biomolecular dynamics. NH is grateful to Huong Vu, Debayan Chakraborty, and Mauro Mugnai for valuable discussions. We thank the Texas Advanced Computing Center at The University of Texas at Austin for providing computational resources. This work was supported in part by grants from the National Science Foundation (CHE 1636424). D.T. also acknowledges additional support from the Collie-Welch Regents Chair (F-0019) administered through the Welch Foundation.

